## Supplemental files for "Impact of blood meals taken on ivermectin-treated livestock on survival and fecundity of the malaria vector *Anopheles coluzzii* under laboratory conditions"

### Appendix S1.

**Preliminary study: efficacy of blood meal taken on treated pigs with therapeutic dose of ivermectin, on the survival of *Anopheles coluzzii***

The methodology was same that the main study. Six (6) pigs were used in the activity (3 control and 3 injected subcutaneously with ivermectin at dose of 0.3 mg/ kg body weight.

A total of 4,155 females were exposed to pigs and 2,170 took blood meal, that is a global engorged rate of 52.27%.

*Survival of An. coluzzii fed on pigs treated with therapeutic dose of ivermectin*

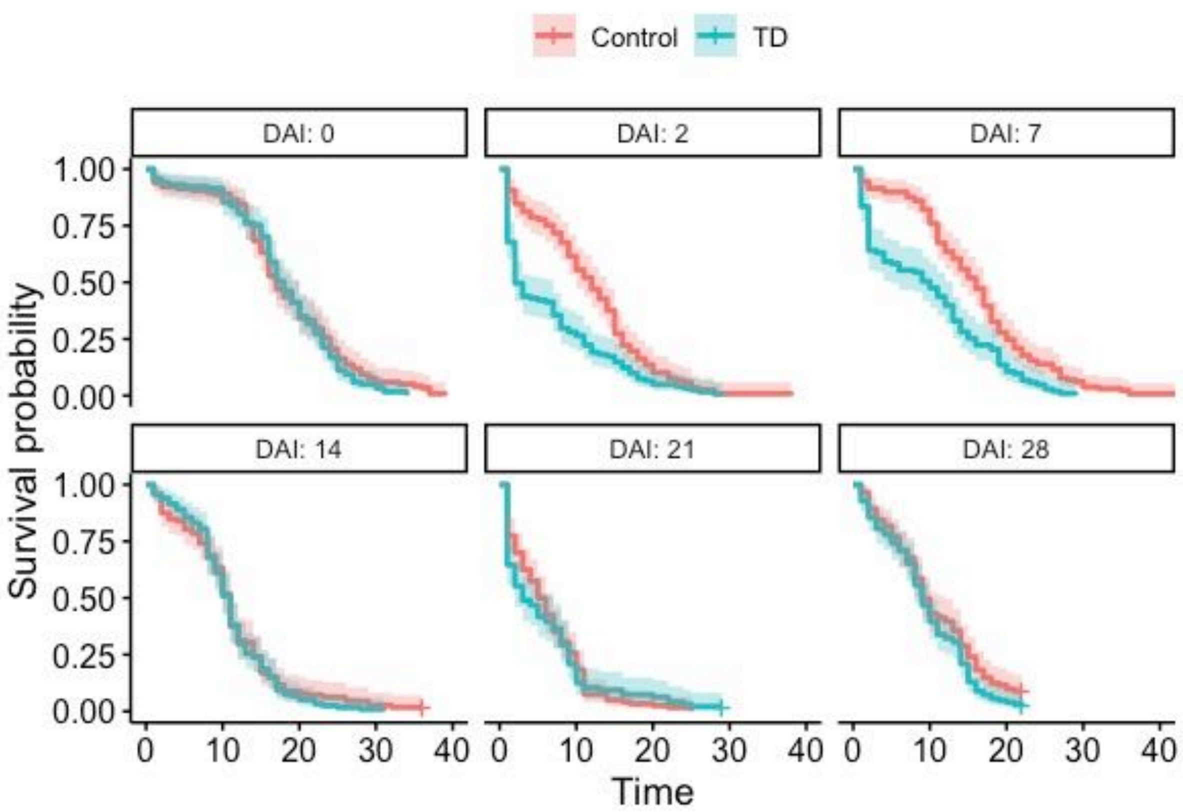


Figure 1. Kaplan-Meier curves showing the survival of mosquitoes fed on pigs from the control and therapeutic dose of ivermectin (TD = 0.3mg/kg) before treatment (DAI = 0) at different Days After Injection (DAI: 2, 7, 14, 21 and 28 days after injection of ivermectin dose.

Before the injection of ivermectin to the pigs during the first experiment, there was no significant effect to individual animal on the anopheles' survival ($X^2_5$ = 6.51; $P$ = 0.26). After the injection of a recommended dose of ivemerctin to pigs, the analysis shows the significant difference of survival due to the effect of the delay post injection ($X^2_5$ = 273.83; $P$ < 0.001), of the ivermectin treatment ($X^2_5$ = 5.59; $P$ = 0.02), and of their interaction ($X^2_5$ = 26.63; $P$ < 0.001) The ivermectin treatment decreased anopheles' survival on day 2 (HR = 1.89, IC [1.43 – 2.50], $P$ < 0.001) and day 7 (HR = 1.92, IC [1.44 – 2.56], $P$ < 0.001).

At 2 DAI, the median survival time of *Anopheles* was 2 days (IC= [2 – 7] days) for the treated group and 12 days (IC= [10 – 14] days) for the control group, representing a decrease of survival of 83.33 % in the treated group. At 7 DAI, the median survival time of *Anopheles* was 10 days (IC= [5 – 13] days) for the treated group whereas it was 16 days (IC= [14 – 17] days) for the control group, representing a decrease of 37.5%.

### <u>Supporting S2 Appendix : Gravidity rate and fecundity in females *Anopheles coluzzii* fed on pig</u>

Before the injection of IVM, gravidity rates of *An. coluzzii* females were different for control and treated groups of pigs (Figure 1). There was a significant lower gravidity rate for females fed on the group of pigs to be treated with the therapeutic dose (OR = 4.00, IC [1.93 – 156], *P* = 0.02) and the double dose group (OR = 4, IC [1.23 – 11], *P* = 0.01) compared to those fed on the control one (Figure 1).

The only significant difference in gravidity rate was observed between *Anopheles coluzzii* females fed on control and treated pigs 7 DAI after injection of the therapeutic dose of ivermectin (OR = 7, IC [1.93 – 22], *P* = 0.004).

Also, the treatment with IVM did not show any effect on *An. coluzzii fecundity,* regardless the treatment dose or the DAI (Figure 2).

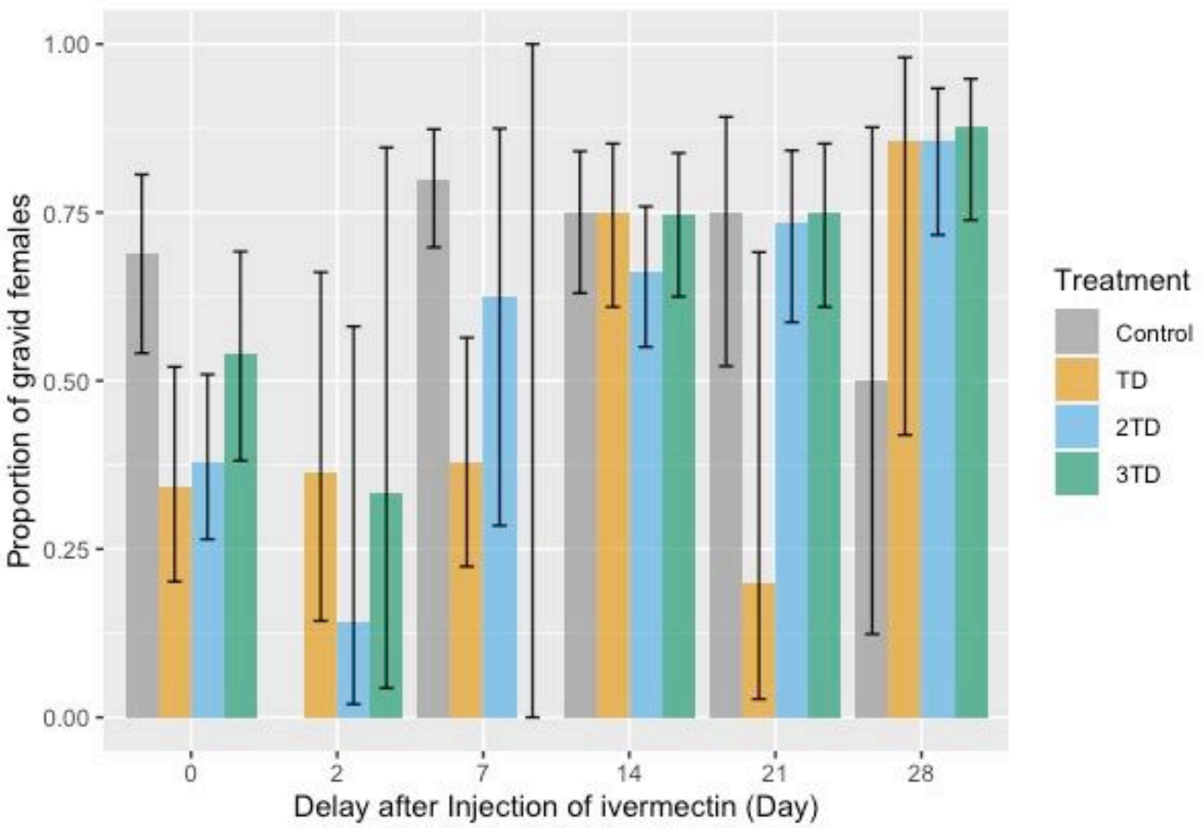


***Figure 1. Proportion of gravid females of Anopheles coluzzii fed on control or treated pigs with different dose of ivermectin (TD: Therzpeutic dose = 0.3 mg/kg; 2TD = 2-fold therapeutic dose; 3TD = 3-fold therapeutic dose). The error bars correspond to the Standard error.***

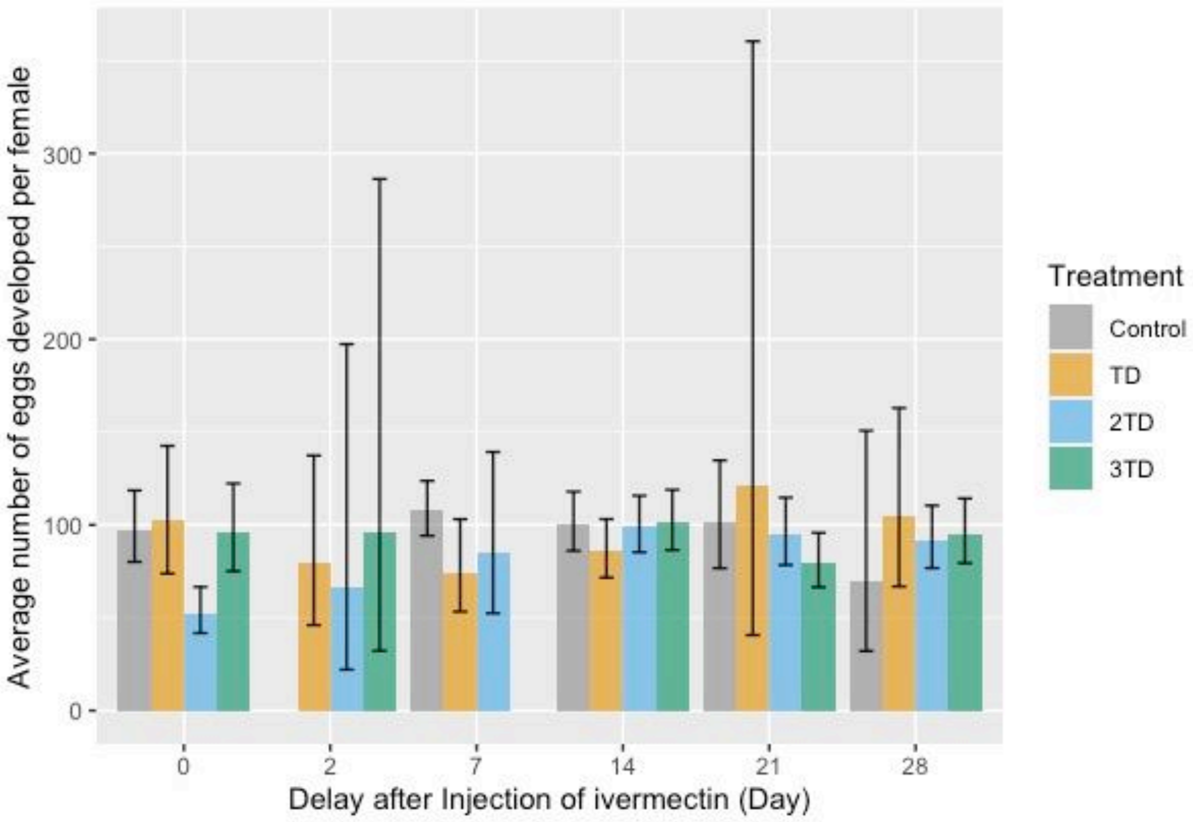


***Figure 2. Average number of eggs developed by gravid females of Anopheles coluzzii fed on control or treated pigs with different dose of ivermectin (TD: Therzpeutic dose = 0.3 mg/kg; 2TD = 2-fold therapeutic dose; 3TD = 3-fold therapeutic dose). The error bars correspond to the Standar error.***

**Supporting S1 Table: Number of exposed, blood-fed and dissected female *Anopheles* according to the hosts species, the treatment and the time elapsed since the injection (DAI)**

| **Hosts species** | **DAI** | **Control** | | | | | **Therapeutic dose** | | | | | **Double therapeutic dose** | | | | | **Triple therapeutic dose** | | | | |
|---|---|---|---|---|---|---|---|---|---|---|---|---|---|---|---|---|---|---|---|---|---|
| | | **T1** | **G1** | **T2** | **G2** | **Diss.** | **T1** | **G1** | **T2** | **G2** | **Diss.** | **T1** | **G1** | **T2** | **G2** | **Diss.** | **T1** | **G1** | **T2** | **G2** | **Diss.** |
| Sheep | | Ivermectin : 0 mg/kg | | | | | Ivermectin at 0.2 mg/kg | | | | | | | | | | | | | | |
| | Before | 472 | 443 | 292 | 121 | 71 | 512 | 471 | 274 | 132 | 99 | - | - | - | - | - | - | - | - | - | - |
| | 2 DAI | 445 | 209 | 138 | 101 | 47 | 555 | 356 | 56 | 34 | 34 | - | - | - | - | - | - | - | - | - | - |
| | 7 DAI | 411 | 205 | 100 | 91 | 60 | 345 | 246 | 90 | 61 | 23 | - | - | - | - | - | - | - | - | - | - |
| | 14 DAI | 196 | 190 | 84 | 72 | 41 | 266 | 173 | 93 | 69 | 55 | - | - | - | - | - | - | - | - | - | - |
| | 21 DAI | 434 | 275 | 148 | 74 | 43 | 372 | 229 | 135 | 73 | 60 | - | - | - | - | - | - | - | - | - | - |
| | 28 DAI | 403 | 269 | 124 | 77 | 54 | 404 | 326 | 182 | 123 | 65 | - | - | - | - | - | - | - | - | - | - |
| ***Sub total (sheep)*** | | ***2361*** | ***1591*** | ***886*** | ***536*** | ***316*** | ***2454*** | ***1801*** | ***830*** | ***492*** | ***336*** | - | - | - | - | - | - | - | - | - | - |
| Goat | | Ivermectin : 0 mg/kg | | | | | Ivermectin at 0.4 mg/kg | | | | | | | | | | | | | | |
| | Before | 643 | 528 | 366 | 209 | 118 | 647 | 590 | 351 | 292 | 117 | - | - | - | - | - | - | - | - | - | - |
| | 2 DAI | 493 | 380 | 159 | 108 | 69 | 502 | 446 | 170 | 137 | 62 | - | - | - | - | - | - | - | - | - | - |
| | 7 DAI | 629 | 524 | 381 | 163 | 118 | 625 | 493 | 297 | 79 | 79 | - | - | - | - | - | - | - | - | - | - |
| | 14 DAI | 547 | 365 | 35 | 89 | 64 | 515 | 297 | 128 | 68 | 68 | - | - | - | - | - | - | - | - | - | - |
| | 21 DAI | 544 | 354 | 145 | 123 | 101 | 514 | 266 | 100 | 86 | 63 | - | - | - | - | - | - | - | - | - | - |
| | 28 DAI | 384 | 258 | 109 | 65 | 44 | 372 | 209 | 58 | 55 | 34 | - | - | - | - | - | - | - | - | - | - |
| ***Sub total (goat)*** | | ***3240*** | ***2409*** | ***1195*** | ***757*** | ***514*** | ***3175*** | ***2301*** | ***1104*** | ***717*** | ***423*** | - | - | - | - | - | - | - | - | - | - |
| Pig | | Ivermectin : 0 mg/kg | | | | | Ivermectin at 0.3 mg/kg | | | | | Ivermectin at 0.6 mg/kg | | | | | Ivermectin at 0.9 mg/kg | | | | |
| | Before | 359 | 165 | 70 | 54 | 45 | 301 | 195 | 90 | 47 | | 373 | 191 | 106 | 76 | 58 | 325 | 171 | 82 | 61 | 37 |
| | 2 DAI | 238 | 80 | 0 | 0 | 0 | 300 | 151 | 34 | 81 | 11 | 223 | 101 | 18 | 17 | 07 | 341 | 125 | 13 | 13 | 03 |
| | 7 DAI | 371 | 284 | 223 | 211 | 80 | 292 | 198 | 67 | 51 | 29 | 343 | 198 | 25 | 17 | 08 | 333 | 258 | 36 | 27 | 09 |
| | 14 DAI | 508 | 268 | 158 | 95 | 64 | 474 | 268 | 166 | 86 | 48 | 442 | 213 | 118 | 81 | 77 | 583 | 412 | 162 | 100 | 63 |
| | 21 DAI | 234 | 130 | 52 | 25 | 20 | 233 | 98 | 17 | 13 | 05 | 267 | 181 | 84 | 52 | 45 | 199 | 137 | 50 | 36 | 48 |
| | 28 DAI | 145 | 81 | 18 | 05 | 04 | 178 | 72 | 11 | 07 | 07 | 230 | 169 | 78 | 45 | 42 | 171 | 146 | 59 | 46 | 41 |
| ***Subtotal (pig)*** | | ***1855*** | ***1008*** | ***521*** | ***390*** | ***213*** | ***1778*** | ***982*** | ***385*** | ***285*** | ***100*** | ***1878*** | ***1053*** | ***429*** | ***288*** | ***237*** | ***1952*** | ***1249*** | ***402*** | ***283*** | ***201*** |
| **General total** | | **7456** | **5008** | **2602** | **1683** | **1043** | **7407** | **5084** | **2319** | **1494** | **859** | **1878** | **1053** | **429** | **288** | **237** | **1952** | **1249** | **402** | **283** | **201** |

Legend: DAI: Day after injection; T1: number of mosquitoes exposed at the first exposition; G1: number of mosquitoes that fed at T1; T2: number of mosquitoes exposed at the second blood-feeding; G2: number of mosquitoes that fed at T2; Diss: number of mosquitoes dissected for fecundity characterization.
